# DeepOrientation: Deep Orientation Estimation of Macromolecules in Cryo-electron tomography

**DOI:** 10.1101/2024.07.12.603241

**Authors:** Noushin Hajarolasvadi, Harold Phelippeau, Antonio Martinez-Sanchez, Robert Brandt, Pierre-Nicolas Suau, Daniel Baum

## Abstract

Orientation estimation of macromolecules in cryo-electron tomography (cryo-ET) images is one of the fundamental steps in applying subtomogram averaging. The standard method in particle picking and orientation estimation is template matching (TM), which is computationally very expensive, with its performance depending linearly on the number of template orientations. In addition to conventional image processing methods like TM, the investigation of crowded cell environments using cryo-ET has also been attempted with deep learning (DL) methods. These attempts were restricted to macromolecule localization and identification while orientation estimation was not addressed due to a lack of a large enough dataset of ground truth annotations suitable for DL. To this end, we first generate a large-scale synthetic dataset of 450 tomograms containing almost 200K samples of two macromolecular structures using the PolNet simulator. Utilizing this synthetic dataset, we address the problem of particle orientation estimation as a regression problem by proposing a DL-based model based on multi-layer perceptron networks and a six-degree-of-freedom orientation representation. The iso-surface visualizations of the averaged subtomograms show that the predicted results by the network are overly similar to that of ground truth. Our work shows that orientation estimation of particles using DL methods is in principle possible provided that ground truth data is available. What remains to be solved is the gap between synthetic and experimental data. The source code is available at https://github.com/noushinha/DeepOrientation.

## 1 Introduction

Recent advances in 3D electron transmission microscopy improved high-resolution imaging of molecular complexes inside native cells using cryo-electron tomography (cryo-ET). In recent years, the throughput of data acquisition in cryo-electron tomography (cryo-ET) has increased thanks to new sample preparation techniques. As a result, spanning the scale of time and quality in current cryo-ET data processing frameworks is inevitable [7]. Cellular cryo-ET attains sub-nanometer resolution inside cells frozen in a near-native state by counting on transmission electron microscopy tilt-series. However, this gain in resolution comes at the cost of a lower signal-to-noise ratio and missing-wedge effect. The standard method for particle identification and orientation estimation is template matching (TM) [1] which suffers from high computational complexity and difficulties in identifying particles with similar structures. In TM, one uses many different orientations of the same template density map for a specific particle and computes the cross-correlation score at every voxel across the whole tomogram. The highest-ranked cross-correlation scores are considered as possible particle locations. In the next step, sub-volumes are extracted at those locations which can be used in turn for other tasks like classification and segmentation.

In addition to conventional image processing methods like TM, the investigation of crowded cell environments using cryo-ET has been attempted with deep learning (DL) methods. Models like DeepFinder [8] and DeePict [3] pick particles and segment tomograms with high accuracy in a reasonable time. However, the lack of annotated data is the major bottleneck to applying DL methods in cryo-ET. Many DL methods with a large number of parameters are not applicable in this field due to the lack of experimental ground truth. An alternative solution to the time-consuming and expensive data acquisition and annotation is training the networks on simulated cryo-ET images. Nonetheless, orientation estimation remains a huge advantage of the TM method over DL-based methods. To the best of our knowledge, none of the DL-based methods developed for processing cryo-ET images addresses the orientation estimation task as a DL-based regression model. It should be highlighted that orientation determination in cryo-ET is computationally very expensive because a comprehensive sampling of 3D rotational space is required. This is mainly because angular space representation based on Euler or quaternions spaces suffer from discontinuity [11] and discontinuous representations are harder to approximate by neural networks [10]. Consequently, converged networks produce large errors at certain rotation angles. We consider a converged network as one where loss value and an evaluation metric like mean-squared error are optimized to minimum value by training.

To address the mentioned problems, we propose a four-layer multi-perceptron (MLP) model that benefits from a continuous representation, called six degrees of freedom (6DoF) [11]. The input to our model is a 3D patch extracted around the particle center and the output is the predicted orientation of the particle in one of the three possible representations namely, Euler, quaternions, or 6DoF.

The main contributions of this work are:

- Contributing to the research community of cryo-ET by publishing large-scale synthetic data; the dataset comprises almost 70K samples in total^1^;
- definition of a unique continuous rotation representation suitable for training a regression neural network on particle orientation estimation.

The rest of this study is organized as follows. We start by reviewing the literature. Then, we discuss data generation, annotation, and introducing 6DoF representation. We continue by proposing a regression model for the problem of macromolecular orientation estimation. Finally, we justify the obtained results using qualitative and quantitative metrics like *r*^2^ score.

## 2 Methods and Materials

As mentioned earlier, investigating the applicability of a dataset is only possible after the last step of the cryo-ET workflow known as tomogram reconstruction. Hence, preparing annotated experimental data is prolonged. A common solution to this issue is generating synthetic data. Current simulators in cryo-ET focus on reproducing distortions from acquired images and reconstructed tomograms. Recently, Martinez et al. proposed a simulator model called PolNet [6] with the capability of generating synthetic cellular environments using several geometric and macromolecular models for low-order structures images of cryo-ET using parametrizable stochastic models. We took advantage of this simulator to generate a large-scale dataset required for training a generalized model. Figure 1 illustrates a cross-section of a synthetic tomogram generated by PolNet.

**Figure 1.**
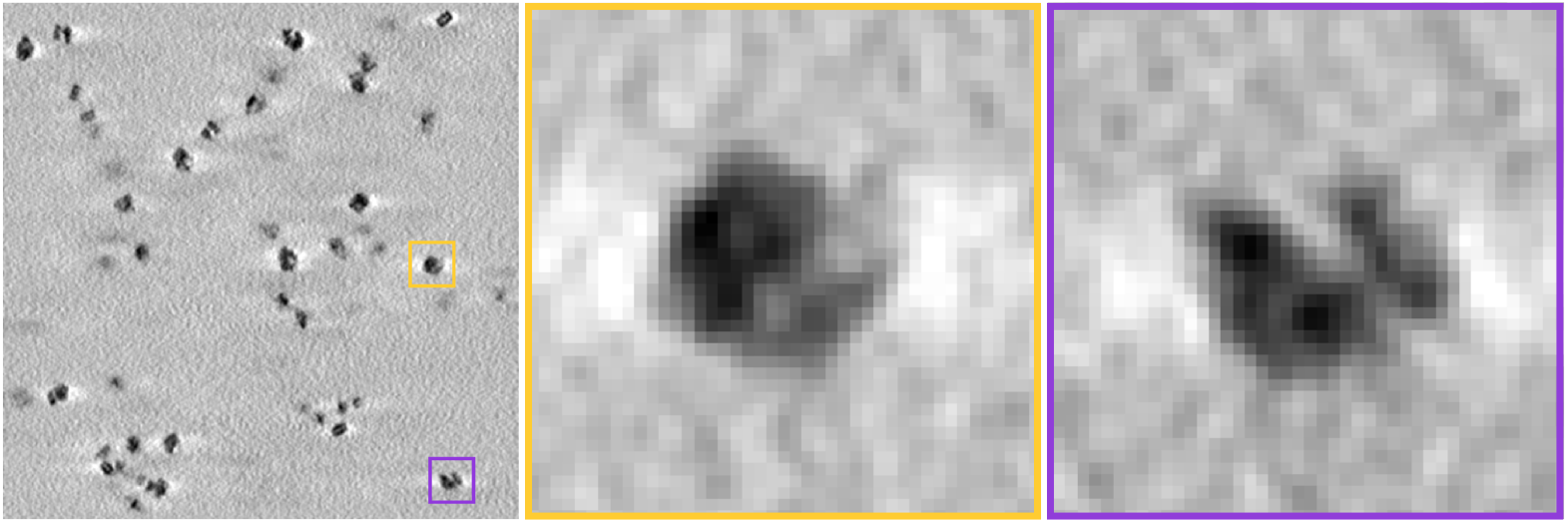
**Left** is the cross-section visualization of particles within a single tomogram generated by PolNet; **middle** and **right** illustrates two magnified regions of the tomogram each containing a ribosome in a specific orientation

### 2.1 Materials

#### 2.1.1 Generating synthetic data with PolNet

Training a generalized model requires a representative and large enough training dataset. We focus on producing rich contextual information by directly generating 3D density maps (synthetic tomograms) containing a variety of protein clusters. For sampling the SO(3) rotational space, we use approximately uniform random distribution as described in [4]. Kernel density estimation of quaternion space generated by PolNet is shown on the left side of Figure 2. The box plot on the right side of the same figure shows that no outlier was generated during the sampling.

**Figure 2.**
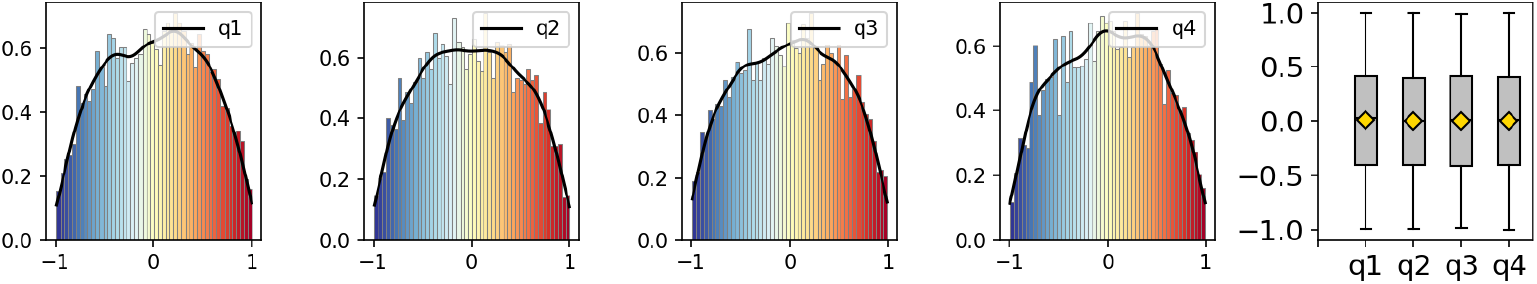
**Left:** We illustrate kernel density estimation of quaternion axis to validate uniform random distribution along each on the ground truth test dataset (4v4r); **right:** the box plots of quaternion sampling distribution in the ground truth test dataset (4v4r) to show zero mean and zero outliers in the distribution

We generated a low-diversity dataset including only two macromolecule types, namely the ribosomal complex (4v4r) and Thermoplasma acidophilum 20S proteasome (3j9i). We select these two specific particles to study the model performance with and without considering the rotational symmetry of the objects. Here, the T20S proteasome 3j9i [5] forms a 700 kDa complex comprising 14 *α* and 14 *β* subunits organized with D7 symmetry and ribosomal complex 4v4r [9] presents the crystal structures of the ribosome from Thermus thermophilus with RF1 and RF2 bound and it is asymmetrical. A comprehensive comparison of the number of samples is given in Table 1.

**Table 1.**
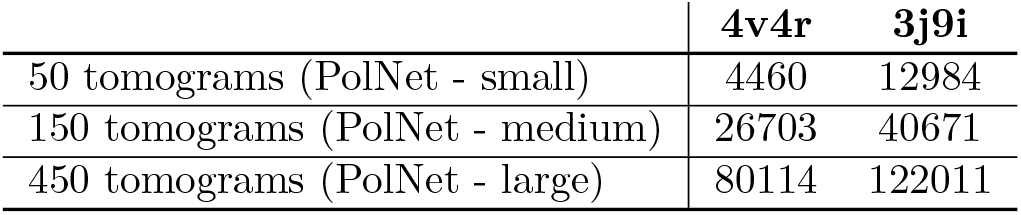
Total number of samples in each dataset per particle.

|  | 4v4r | 3j9i |
| --- | --- | --- |
| 50 tomograms (PolNet - small) | 4460 | 12984 |
| 150 tomograms (PolNet - medium) | 26703 | 40671 |
| 450 tomograms (PolNet - large) | 80114 | 122011 |

We also study the effect of dataset size on our model considering the overfitting and underfitting problem by generating three datasets having different numbers of tomograms that we call small, medium, and large. The small dataset includes only 50 tomograms with ∼ 4*K* samples of the ribosome and ∼ 13*K* samples of the proteasome. The small dataset has 40 tomograms of size 500 × 500 × 250 and 10 tomograms of size 1000 × 1000 × 250 voxels with a 10 Å voxel size. The medium dataset comprises 150 tomograms with ∼ 27*K* and ∼ 41*K* samples for ribosome and proteasome, respectively. Later, due to the overfitting problem, we extended the medium-scale dataset by generating 300 additional tomograms to have almost 80K ribosome samples and 122K proteasome samples distributed across 450 tomograms. All tomograms of medium and large datasets are of size 500 × 500 × 250 voxels and 10 Å voxel size. Figure 3 illustrates the effect of data sample size on model generalization using small, medium, and large datasets. Further results on these experiments are reported in Section 3.

**Figure 3.**
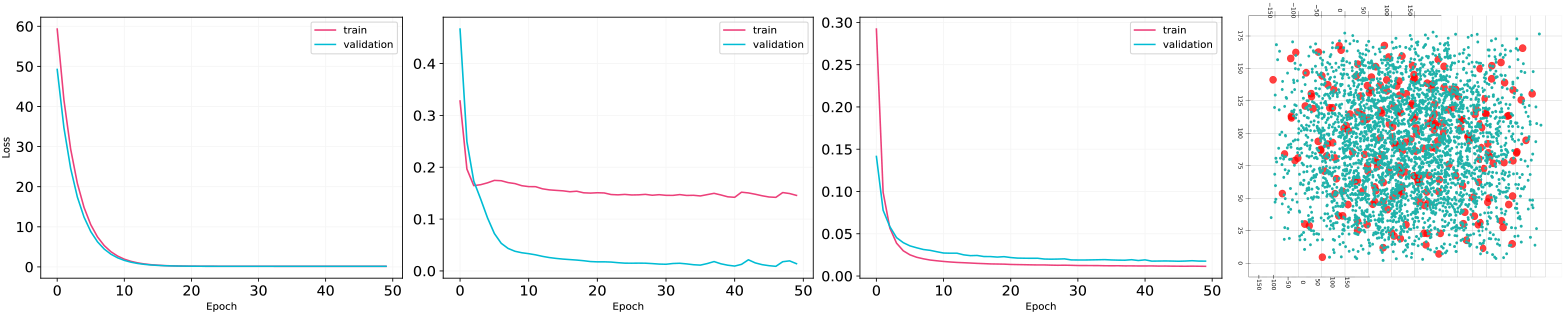
**Left:** Training plots using a small, medium, and large size dataset with 4K, 27K, and 80K ribosome samples that underfitted, overfitted, and optimally fitted to the data after training; underfitting and overfitting problems are solved with increasing complexity of the model and using a larger dataset; **right:** Projected distribution of samples based on two axes of the Euler angles for small (blue) and large (pink) datasets

### 2.2 Methodology

Our approach is based on the work by Zhou et al. [11]. They study the continuity of orientation representation in the neural network context. We first introduce our general notation and we continue by establishing the 6DoF representation.

#### Notation

We denote a matrix as *M* and so *M_ij_* represents the (*i, j*) element of the matrix. The term *SO*(*n*) denotes the space of *n*-dimensional rotations which is a special orthogonal group defined on the set of *n* × *n* real matrices with multiplication operation and holding properties *M^T^ M* = *MM^T^* = *I* and *det*(*M*) = 1 true. An *n*-dimensional unit sphere is then denoted as *S^n^* = *{x* ∈ ℝ^*n*+1^ : ||*x*|| = 1}.

#### Discontinuous representation

We meet discontinuity problems for 2D rotations which can be extended to *n*D rotations. Under given definitions, any 2D rotation *M* ∈ *SO*(2) can be expressed as:

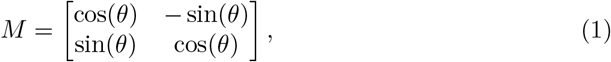

where *θ* ∈ ℛ is an angle, and ℛ can be [0, 2*π*]. In such case, a mapping *g* from original space *SO*(2) to the angular representation space ℛ is discontinuous because the limit of *g* at the identity matrix that represents no rotation, is undefined. In particular, the limit of such a mapping in one direction gives an angle of 0 while in the other direction, it gives 2*π*. This phenomenon is shown in Figure 4, left.

**Figure 4.**
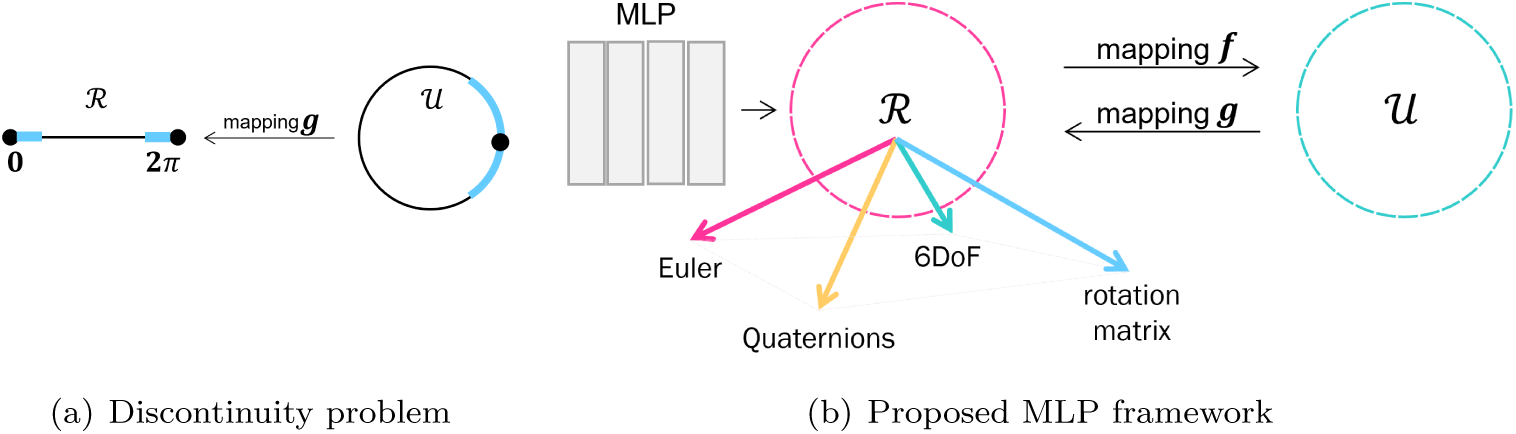
**Left:** discontinuity problem in a 2D representation space; **right**: the block diagram of the proposed multi-layer perceptron model with 4 layers

#### Continuous representation

Let ℛ indicate a *representation space*, i.e. a subset of real vector space where the Euclidean distance is defined. Also, let 𝒰 be a compact topological space that we call *original space*. Then, based on the approach by Zhou et al. [11], we consider a neural network that employs an intermediate representation in ℛ. These representations can be mapped into the original space 𝒰, if we define the mapping to the original space as *f* : ℛ → 𝒰 and mapping to the representation space *g* : 𝒰 → ℛ. We call (*f, g*) a representation if ∀*u* ∈ 𝒰 *, f* (*g*(*u*)) = *u*. We say the representation is continuous if g is continuous.

#### Continuity in Neural Network context

Given an input to the network, it should output a representation in ℛ, then pass it through mapping *f* to get an element of the original space 𝒰 . The right side of Figure 4 illustrates this concept. Here, mapping *f* and *g* helps to solve the discontinuity problem. Based on this observation, we propose to perform an orthogonalization like Gram-Schmidt in the representation space itself. Let original space be 𝒰= *SO*(*n*), and the representation space ℛ = *ℝ*^*n*×(*n*−1)^ \ *D* where *D* is the set of elements that the following Gram-Schmidt process cannot map to *SO*(*n*). Then the mapping *g_GS_* to ℛ can be expressed as omitting the last column of the rotation matrix:

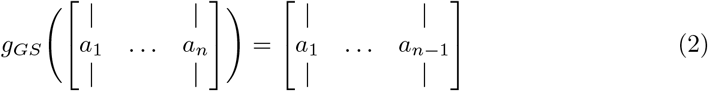

Here, *a_i_* for *i* = 1, 2*, *n*, n* are column vectors, *g_GS_*(𝒰) is the set of all orthonormal k-frames in ℝ*^n^*. Now, for mapping *f_GS_* to the original space, a Gram-Schmidt-like [11] process can be defined:

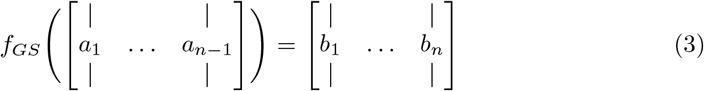

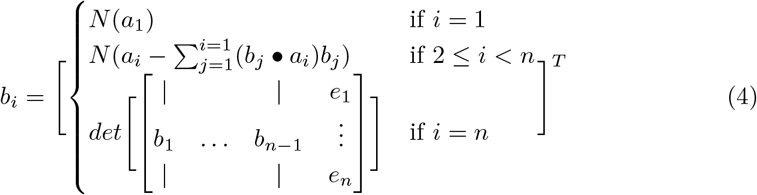

where *N* (·) is the normalization function, and *e*_1_, *n*, *e*_*n*_ are the *n* basis vectors of Euclidean space. The aforementioned expressions are the general case of continuity. Having 3 × 3 rotation matrices, the representation with six-degree-of-freedom (6DoF) can be obtained by dropping the third column of the rotation matrix using equation (3). Later, by applying equation (4) the original rotation matrix can be reconstructed.

#### DeepOrientation

We use an MLP network to learn the proposed 6DoF representation space. We test 6DoF against other representation spaces, namely, Euler and quaternions. The model consists of four dense layers where each layer has 128 nodes and uses ReLU activation. We apply Huber loss between the ground truth and the corresponding estimated output. The network is trained for 50 epochs without early stopping criteria. In addition to mean-squared-error (MSE) and mean-absolute-error (MAE), we compute two custom metrics. First, the *r*^2^ score is a statistical measure used to determine the proportion of variance in a dependent variable that can be predicted or explained by an independent variable. It has a value between 0 and 1 and the higher it is the better the performance. Second, cosine similarity angular error is defined as the minimum difference between two vectors *a* and *b*, formulated as arcoss 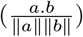.

## 3 Results

The code was implemented in Keras and we used Keras tuner for hyperparameter tuning of the network. Specifically, we applied hyperband, random search, and Bayesian optimization on a small set of data to find the optimal number of nodes, number of layers, batch size, learning rate, and dropout for the MLP network architecture. We should note that hyperband tuning worked the best in our case. We fixed our network and then used this set of best parameters for all the following experiments unless it is mentioned otherwise. As best parameters, we determined 4 layers, 128 nodes per layer, 1e-04 learning rate, batch size 32, centered patches of size 40^3^, 50 epochs, Adam optimizer, and Huber loss. We always used 85% of the training data for training and 15% for validation. Data was normalized (zero mean, unit variance) and shuffled.

### 3.1 Performance Analysis on synthetic data

#### Dataset size

As mentioned earlier, we performed three experiments to avoid overfitting problems. Table 2 shows the improvement of *r*^2^ score and MSE as the dataset size is increased.

**Table 2.** Comparison of statistical metrics for different dataset sizes (4v4r)

| | | $r^2$ | MSE | error |
| --- | --- | --- | --- | --- |
| Small | train | 0.8134 | 0.0621 | $20^\circ \pm 28$ |
| | test | -0.0162 | 0.3379 | $84^\circ \pm 23$ |
| Medium | train | 0.9633 | 0.0122 | $7^\circ \pm 6$ |
| | test | 0.9582 | 0.0139 | $15^\circ \pm 14$ |
| Large | train | 0.9785 | 0.0071 | $6^\circ \pm 4$ |
| | test | 0.9585 | 0.0138 | $8^\circ \pm 9$ |

The angular error described in Section 2.2 decreased as the number of samples increased. Figure 5 represents the model convergence based on the MSE metric. Clearly, the small dataset has the largest mean. Outliers in medium and large data belong to the first few epochs of training. Additionally, Figure 6(b) shows a histogram chart of the number of correctly estimated orientations within the test dataset of 4v4r samples. It is important to note that we keep the test data fixed for fair comparison purposes during the experiments on medium and large data.

**Figure 5.**
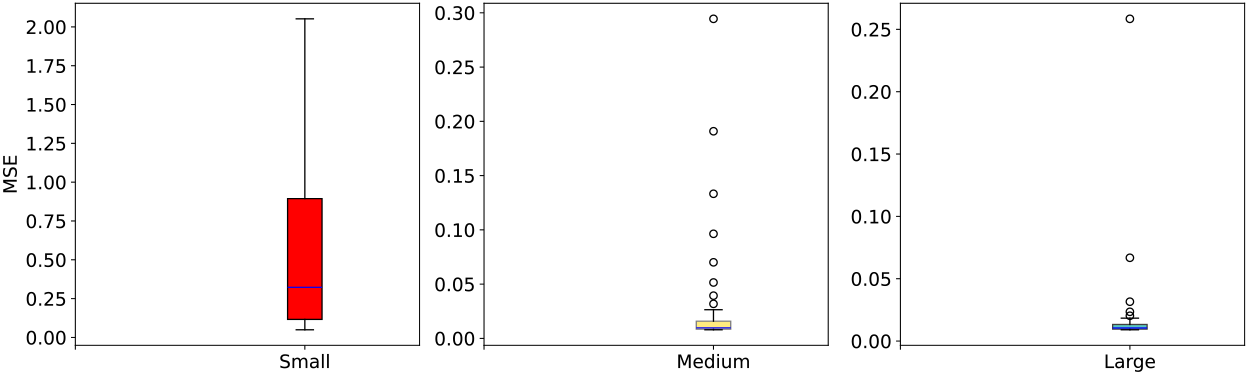
Illustration of MSE convergence for small, medium, and large datasets (4v4r)

**Figure 6.**
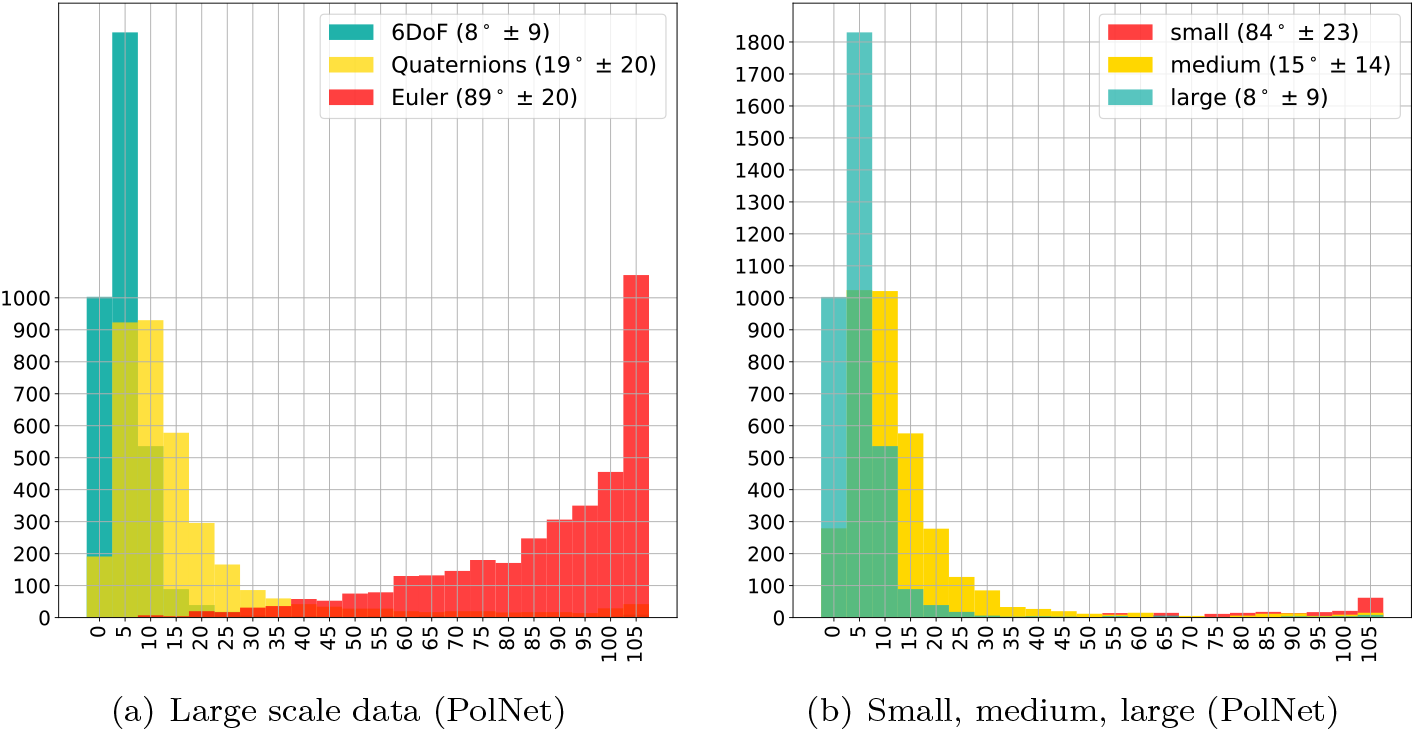
Histogram chart of the angle differences with a 5^°^ bin width for 4v4r; a) the network estimated 0.96% of particles accurately using 6DoF, b) using a larger dataset increased number of correct estimations

As a result, the test dataset of 4v4r particles for these two datasets has 3575 samples that are distributed among 20 tomograms. The small dataset of 4v4r includes 228 samples in a single tomogram.

#### Different orientation representations

Here, we analyze the performance of DeepOrientation for the aforementioned orientation representations. We train the network using Euler, quaternions, and 6DoF representations of 4v4r and 3j9i separately. Table 3 compares these results statistically for 4v4r particles.

**Table 3.**
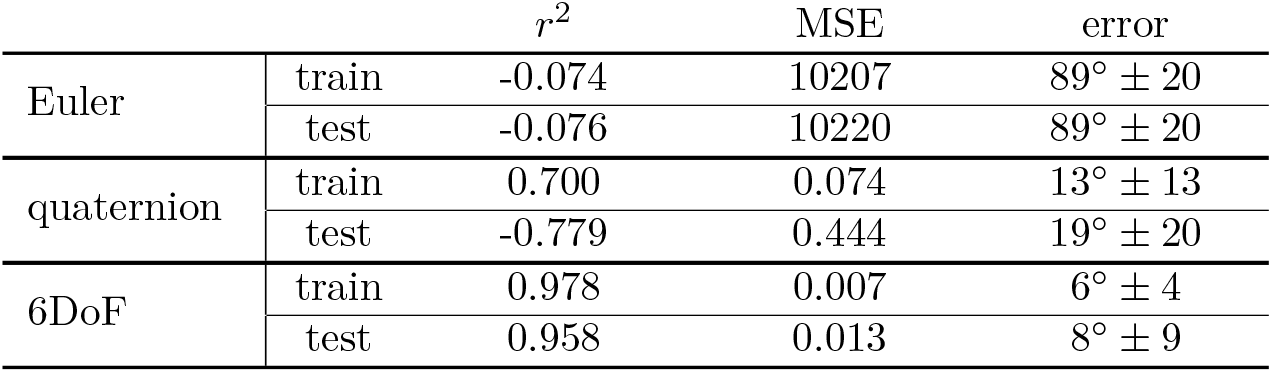
Comparison of statistical metrics for training the network on ∼80K samples of 4v4r using Euler, quaternions, 6DoF representations.

| | | $r^2$ | MSE | error |
| --- | --- | --- | --- | --- |
| Euler | train | -0.074 | 10207 | $89^\circ \pm 20$ |
| | test | -0.076 | 10220 | $89^\circ \pm 20$ |
| quaternion | train | 0.700 | 0.074 | $13^\circ \pm 13$ |
| | test | -0.779 | 0.444 | $19^\circ \pm 20$ |
| 6DoF | train | 0.978 | 0.007 | $6^\circ \pm 4$ |
| | test | 0.958 | 0.013 | $8^\circ \pm 9$ |

Crowther criterion [2] is known as the minimum angle difference that can be measured on a discrete representation of a specific macromolecule at a given resolution and radius, i.e. it defines the minimum rotation sampling at a given resolution and can be defined as 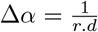 where *d* and *r* are the particle diameter and resolution of the tomogram in Å and Å^−1^, respectively, and the output Δ*α* becomes the angular increment in radians. By such calculation, the value of Δ*α* is equal to 0.038 radians or 2.17^°^ for 4v4r particle and 0.066 radians or 3.78^°^ for 3j9i. That is the smallest rotation one can attain under this configuration.

To plot the histogram charts, we allow a 10-step error tolerance of the desired threshold, ∼ 20^°^. Figure 6(a) compares the number of correctly estimated orientations with an angle difference of 5^°^ bin width using the aforementioned threshold based on Crowther criterion for the three orientation representations and three different dataset sizes of 4v4r particle. As the histogram charts show in Figure 6(a), the network estimated 3459 particles out of 3575 correctly when trained using 6DoF representation, missing only 116 particles. This number decreases to 2623 and 14 for quaternions and Euler, respectively. In the case of the small dataset, the network misses estimating almost all test data samples in the small dataset using the best representation (6DoF) with only 3 out of 228 correct estimations. Unfortunately, measuring the angle diffence is not meaningful for 3j9i due to the rotational symmetry of the particle. As a result, we use an alternative metric that works regardless of the rotational symmetry, i.e. averaged sub-tomogram. The test dataset of 3j9i includes 5418 samples. We compute an averaged subvolume free of distortions (noise and missing wedge) from the original tomograms by aligning all instances of the 3j9i particle according to the predicted rotation. Consequently, the higher the similarity between the averaged and original models, the higher the accuracy for the predicted rotations. Later, we visualize the iso-surface of the computed sub-tomogram and we align it with the corresponding molecular structure. Our experimental results show that the predicted orientations can successfully reconstruct the surface of both particles. These qualitative results are depicted in Figure 8.

**Figure 7.**
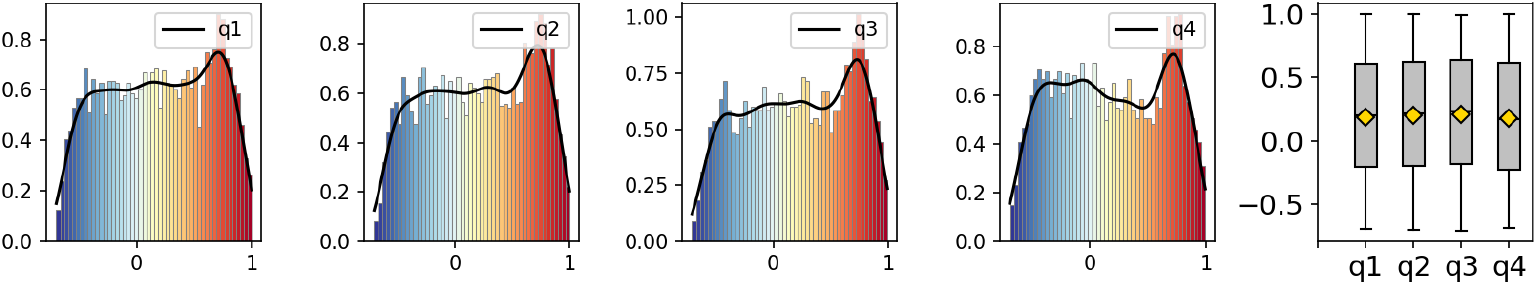
**Light:** We illustrate kernel density estimation of quaternion axis to validate uniform random distribution along each on predictions of test dataset (4v4r); **right:** the box plots of quaternion sampling distribution in predictions of test dataset (4v4r) to show almost zero mean and no outlier in the predicted distribution

**Figure 8.**
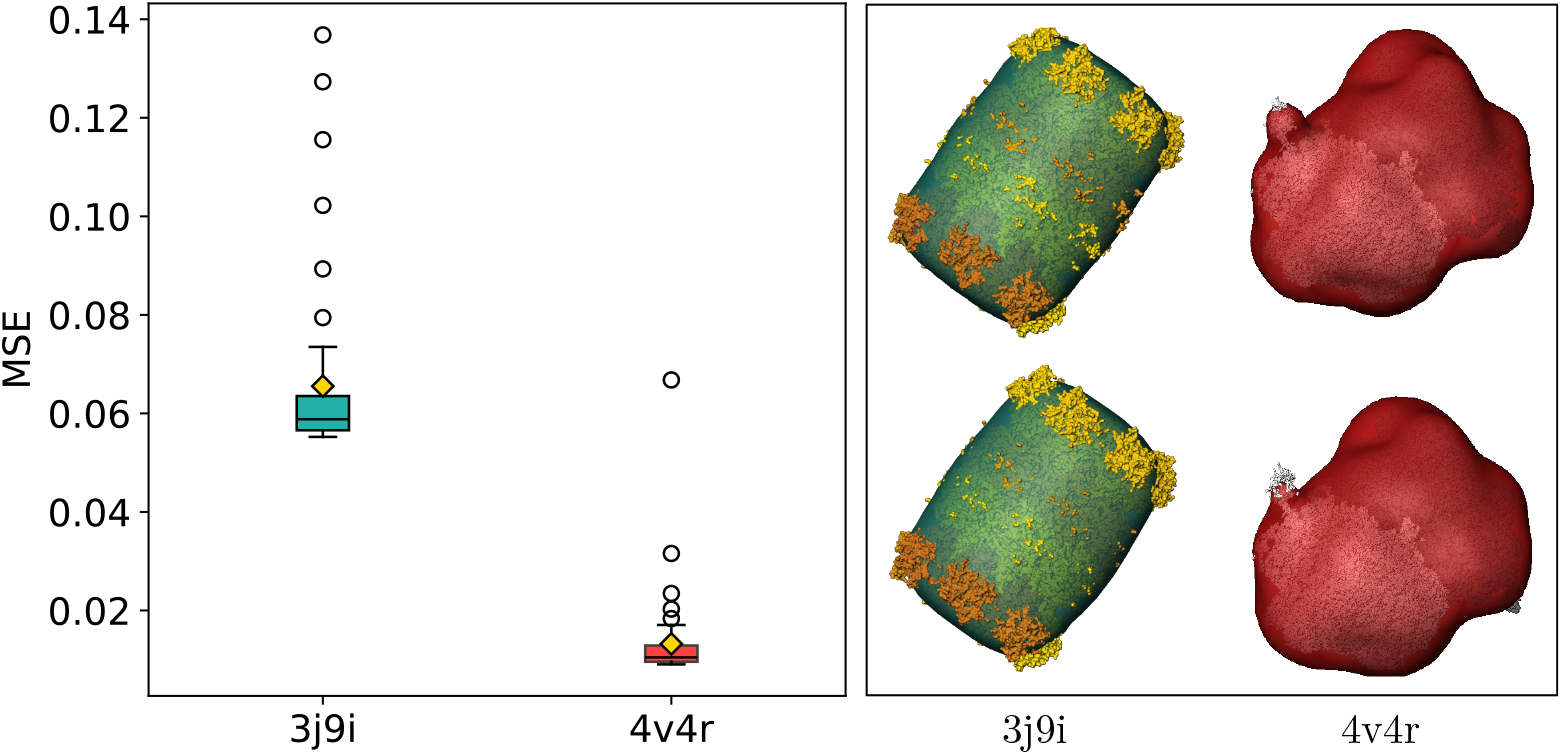
**Left:** Network convergence based on MSE metric for 3j9i and 4v4r particles, The network shows slightly better performance on asymmetrical particles; **right:** top row shows the iso-surfaces computed from 5418 and 3575 test samples of the ground truth for 3j9i and 4v4r; bottom row is the iso-surface computed using the estimated orientations by the network on the same test dataset.

Even though *r*^2^ is a fitting measure for regression models, it has limitations. For instance, small *r*^2^ values do not always show a low performance, as is the case for 3j9i particles in our study. Despite the lower values of *r*^2^ and the angular difference in comparison with 4v4r as an asymmetrical particle, the computed averaged sub-tomograms using the predicted orientations are close to that of ground truth. Our results show that the network indeed shows a good performance on both symmetrical and asymmetrical particles in terms of the averaged sub-tomogram metric and network convergence.

## 4 Future Work

In this paper, we developed a deep learning-based method to address the orientation estimation of macromolecules in cryo-ET images. We consider a four-layer multi-perceptron network and feed pairs of sub-tomograms with corresponding orientation representation known as 6DoF to the network. The network performance is evaluated for different dataset sizes with one symmetrical and one asymmetrical particle. Our large dataset includes almost 200K samples in total. The performance of the network is evaluated using MSE as well as averaged sub-tomograms and iso-surface computation metrics. Our future work aims to compare these results with the baseline conventional methods, namely template-matching. To this end, we aim to design a hybrid network where both the position and orientation of the macromolecules are learned using an end-to-end model.

## Acknowledgments

This work was supported by the German Ministry for Education and Research (BMBF) within the Forschungscampus MODAL (project grant 3FO18501). A.M-S. is supported by the Ramon y Cajal Program of the Spanish State Research Agency (AEI) through the MICIU/AEI/10.13039/501100011033 and the European Union NextGenerationEU/PRTR under Grant RYC2021-032626-I and University of Murcia through the Attract-RYC 2023 Program.

## Footnotes

1 https://www.zib.de/ext-data/PolNet_Medium_Size_Dataset_4v4r_and_3j9i.zip

## References

1. C. Best, S. Nickell, and W. Baumeister. Localization of protein complexes by pattern recognition. Methods in cell biology, 79:615–638, 2007.

2. M. L. Chaillet, G. van der Schot, I. Gubins, S. Roet, R. C. Veltkamp, and F. Förster. Extensive angular sampling enables the sensitive localization of macromolecules in electron tomograms. International Journal of Molecular Sciences, 24(17):13375, 2023.

3. I. de Teresa-Trueba, S. K. Goetz, A. Mattausch, F. Stojanovska, C. E. Zimmerli, M. Toro-Nahuelpan, D. W. Cheng, F. Tollervey, C. Pape, M. Beck, et al. Convolutional networks for supervised mining of molecular patterns within cellular context. Nature Methods, 20(2):284–294, 2023.

4. J. Kuffner. Effective sampling and distance metrics for 3d rigid body path planning. In IEEE International Conference on Robotics and Automation, 2004. Proceedings. ICRA ‘04. 2004, volume 4, pages 3993–3998 Vol.4, 2004.

5. X. Li, P. Mooney, S. Zheng, C. R. Booth, M. B. Braunfeld, S. Gubbens, D. A. Agard, and Y. Cheng. Electron counting and beam-induced motion correction enable near-atomic-resolution single-particle cryo-em. Nature methods, 10(6):584–590, 2013.

6. A. Martinez-Sanchez, L. Lamm, M. Jasnin, and H. Phelippeau. Simulating the cellular context in synthetic datasets for cryo-electron tomography. bioRxiv, pages 2023–05, 2023.

7. C. L. McCafferty, S. Klumpe, R. E. Amaro, W. Kukulski, L. Collinson, and B. D. Engel. Integrating cellular electron microscopy with multimodal data to explore biology across space and time. Cell, 187(3):563–584, 2024.

8. E. Moebel, A. Martinez-Sanchez, L. Lamm, R. D. Righetto, W. Wietrzynski, S. Albert, D. Larivière, E. Fourmentin, S. Pfeffer, J. Ortiz, et al. Deep learning improves macromolecule identification in 3d cellular cryo-electron tomograms. Nature Methods, 18(11):1386–1394, 2021.

9. S. Petry, D. E. Brodersen, F. V. Murphy, C. M. Dunham, M. Selmer, M. J. Tarry, A. C. Kelley, and V. Ramakrishnan. Crystal structures of the ribosome in complex with release factors rf1 and rf2 bound to a cognate stop codon. Cell, 123(7):1255–1266, 2005.

10. Z.-B. Xu and F.-L. Cao. Simultaneous lp-approximation order for neural networks. Neural Networks, 18(7):914–923, 2005.

11. Y. Zhou, C. Barnes, J. Lu, J. Yang, and H. Li. On the continuity of rotation representations in neural networks. In Proceedings of the IEEE/CVF conference on computer vision and pattern recognition, pages 5745–5753, 2019.

